# GDmicro: classifying host disease status with GCN and Deep adaptation network based on the human gut microbiome data

**DOI:** 10.1101/2023.06.12.544696

**Authors:** Herui Liao, Jiayu Shang, Yanni Sun

**Affiliations:** Department of Electrical Engineering, City University of Hong Kong, Kowloon, Hong Kong (SAR), China

## Abstract

**Motivation:** With advances in metagenomic sequencing technologies, there are accumulating studies revealing the associations between the human gut microbiome and some human diseases. These associations shed light on using gut microbiome data to distinguish case and control samples of a specific disease, which is also called host disease status classification. Importantly, using learning-based models to distinguish the disease and control samples is expected to identify important biomarkers more accurately than abundance-based statistical analysis. However, available tools have not fully addressed two challenges associated with this task: limited labeled microbiome data and decreased accuracy in cross-studies. The confounding factors such as the diet, technical biases in sample collection/sequencing across different studies/cohorts often jeopardize the generalization of the learning model.

**Results:** To address these challenges, we develop a new tool GDmicro, which combines semi-supervised learning and domain adaptation to achieve a more generalized model using limited labeled samples. We evaluated GDmicro on human gut microbiome data from 10 cohorts covering 5 different diseases. The results show that GDmicro has better performance and robustness than state-of-the-art tools. In particular, it improves the AUC from 0.783 to 0.949 in identifying inflammatory bowel disease. Furthermore, GDmicro can identify potential biomarkers with greater accuracy than abundance-based statistical analysis methods. It also reveals the contribution of these biomarkers to the host’s disease status.

**Availability and implementation:** https://github.com/liaoherui/GDmicro

**Supplementary information:** Supplementary data are available at XXX online

## Introduction

In recent years, many studies have shown strong associations between the human gut microbiome and several human diseases [1, 2, 3]. For example, a meta-analysis of large-scale metagenomic samples shows that dozens of specific bacteria are enriched in colorectal cancer (CRC) patients across different countries [4]. Another microbiome-related study found a reduced complexity of the bacterial phylum *Firmicutes* in inflammatory bowel disease (IBD) patients [5]. With the in-depth study of *Firmicutes*, anti-inflammatory properties of many species under this phylum have been revealed, implying their potential utilities in promoting gut health. These observations indicate that the composition of gut microbes may provide important information for distinguishing case and control samples of a particular disease. Given the accumulating evidence on the associations between diseases and the human gut microbiome, there is a need to develop a more accurate microbiome-based host disease status classification model. Such a model has the potential to identify more informative biomarkers for downstream analysis than abundance-based statistical analysis [6]. The goal of this study is to create a more accurate host disease status classification model using data from the human gut microbiome and incorporating advanced functions for biomarker discovery.

Many computational methods have been developed to classify host disease status based on the human gut microbiome data. The abundance of gut microbes is a major feature used by these tools [6]. Given the gut microbial composition abundance data, these methods apply traditional machine learning or deep learning methods to distinguish case and control samples of a specific disease. The tools utilizing traditional machine learning include MetAML [7] and SIAMCAT [8]. MetAML takes gut microbial compositional data as input and then applies random forest and support vector machines to classify host disease status. To improve the classification performance, it combines k-fold cross-validation and the grid search strategy to tune the best hyperparameters for models. SIAMCAT [8] is a toolbox that relies on various regression models, such as ridge regression, to classify host disease status using gut microbial compositional data.

In addition to traditional machine learning methods, deep learning models have shown promising results in classifying host disease status [9, 10]. One relevant work is DeepMicro [9], which applies an autoencoder model to learn the deep representation of input gut microbial compositional features. Then, the multi-layer perceptron (MLP) model is employed to classify host disease status with the learned latent features. PopPhy-CNN [10] is another popular deep-learning-based tool. It takes gut microbial compositional data and a phylogenetic tree as input and utilizes a novel convolutional neural network (CNN) learning framework for host disease status classification. Despite the promising results, existing tools have not addressed two challenges well. The first challenge is the limited number of labeled data. Although there has been rapid growth in microbiome data collection, a large number of samples still lack detailed metadata annotations. According to a recent study, only 7.8% of the 444,829 human microbiome samples had explicit information on host disease status [11], partly because of the high cost of obtaining labels for microbiome data [12]. As a result, limited labeled data pose challenges for most learning models. Second, many available methods ignore the domain discrepancy problem. For example, in our k-fold cross-validation experiment on CRC, DeepMicro achieves a 0.803 AUC on the CRC-FR dataset. However, when applied to CRC-US dataset in the cross-study experiment, its performance drops to 0.609. Specifically, data from different studies have many differences due to confounding factors such as region, ethnicity, and diet, which can lead to changes in the gut microbiome [4, 13]. Thus, the domain difference between training and test data can greatly affect the robustness of the classification model, ultimately leading to poor performance in real applications.

### Overview of our method

Given the limited labeled data and the rapid accumulation of unlabeled data, we formulate the host disease status classification problem as a semi-supervised learning problem, which uses both labeled and unlabeled data for feature learning. Here, we present GDmicro, a graph convolutional network (GCN) based model to utilize the information from both labeled and unlabeled samples for learning and classification. First, an inter-host microbiome similarity graph is built using the gut microbial compositional data. In our graph, nodes represent the compositional abundance features of the hosts’ microbiome (Fig. 1 I), and edges represent the similarity of learned latent features between two hosts’ microbiome (Fig. 1 II-III). Then, GCN can take this graph as input and incorporates the structural and node abundance features for disease status classification. Because both labeled and unlabeled samples can be integrated into the graph, graph convolution in GCN can utilize local topology for feature passing between two types of samples. Second, to overcome the domain discrepancy problem, we apply a deep adaptation network to learn transferable latent features from the microbial compositional matrix across different domains (Fig. 1 II). We validated GDmicro on 10 cohorts covering 5 diseases and compared GDmicro with alternative tools for disease status classification. The results show that GDmicro has consistently higher AUC than other methods. In addition, GDmicro allows users to detect the most important species to disease status classification and explore how these species affect the hosts’ disease status, which provides important information for biomarker discovery.

**Fig. 1.**
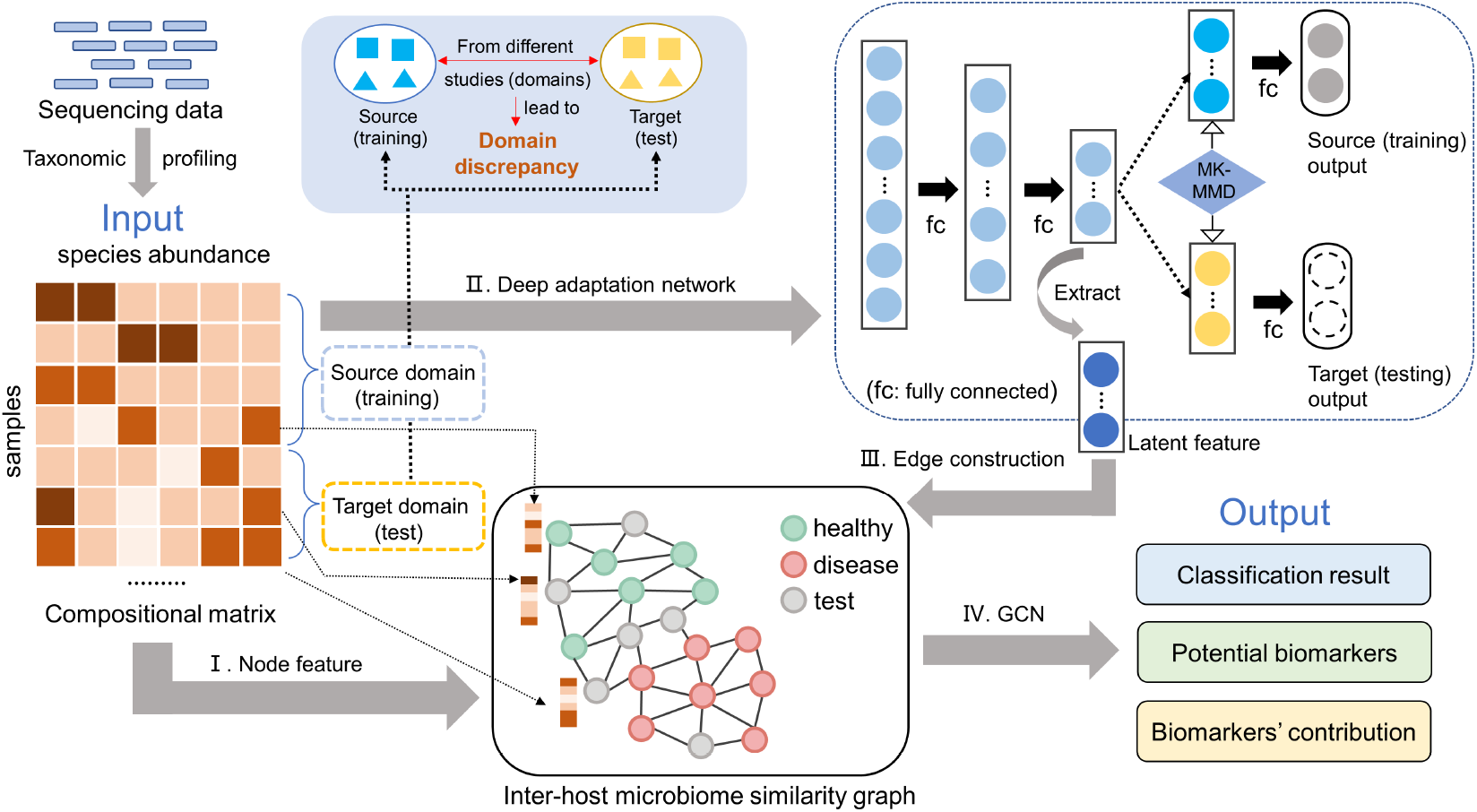
The overall workflow of GDmicro. I. Representing nodes with species abundance vectors in the inter-host microbiome similarity graph. II. Applying the deep adaptation network to learn transferable latent features. The deep adaptation network reduces the domain discrepancy between training and test data. III. Constructing edges of the inter-host microbiome similarity graph with the Euclidean distance of learned latent features from compositional data. IV. Training a GCN model to classify disease status for test samples.

## Methods

We choose GCN as the semi-supervised learning engine for our host disease status classification problem, which uses both labeled and unlabeled data for feature learning. GCN has had several successful applications in identifying gene-gene interaction, disease-drug relationship, and interactions between phages and bacteria [14, 15, 16]. In the host disease status classification problem, GCN has two distinct advantages. First, it can utilize information from unlabeled samples during graph convolution. Second, the GCN model systematically integrates information from the nodes and edges, which represent the abundance distribution of species and their similarities between hosts. Then, we apply the deep adaptation network to mitigate the impact of domain-specific confounding factors on feature learning, leading to a more robust graph for microbiome data from different studies.

In the following sections, we will first introduce how we encode and construct the inter-host microbiome similarity graph with the gut microbiome data and deep adaptation network. Then, we will describe the GCN model optimized for disease identification and its application in biomarker discovery.

### Inter-host microbiome similarity graph *G*

Increasing studies show that people with sclerosis, obesity, and inflammatory bowel diseases have similar gut microbial compositional abundance [17, 18, 19]. One recent work constructed a patient relationship graph based on the similarity of human multi-omics data for disease diagnosis [20]. Inspired by these studies, we construct an inter-host microbiome similarity graph *G* to incorporate microbial compositional similarity and compositional abundance features. The major components of the graph construction are sketched in Fig. 1 (I-III). Each node in the graph represents a human microbial sample, and the edges represent the similarity between the microbiome data. The node features are the compositional abundance vectors of the samples. Because the compositional abundance data might contain skewness or bias in some features, we apply log10-transformation and z-score normalization on the features rather than using the raw data. As a result, each node is encoded with the normalized species abundance vectors.

As mentioned above, similar gut microbial compositional data can indicate similar human disease status. Thus, we use microbial compositional similarity to define the edge between two nodes. Because there is no clear cutoff to determine whether two samples are significantly similar, we employ the *k*-nearest neighbor algorithm to generate the topological structure. Specifically, a node will be connected with its *k* closest nodes in *G*. Below we will detail how to compute the compositional similarity between different samples (Fig. 1 II-III).

### Using deep adaptation network to learn transferable features

To learn the most relevant features for edge construction from different studies, we applied a deep adaptation network to learn transferable latent features from compositional abundance matrices of training and test datasets.

The deep adaptation network is initially designed to solve the domain adaptation problem in the field of image processing [21]. The main idea behind this is to utilize the multiple kernel variants of maximum mean discrepancies (MK-MMD) [22] to measure the difference between the source and the target domain and minimize the domain discrepancies during training. To apply MK-MMD to microbiome data, we added an MK-MMD-based adaptation regularizer to the loss function of a multi-layer fully connected network (Fig. 1 II). Denote *x* as the compositional abundance vector of one sample, *y* as the disease status label of the sample, 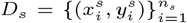 as the source domain with *n*_*s*_ labeled samples and 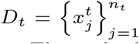 as the target domain with *n*_*t*_ unlabeled samples. Then, the loss function of the network can be defined as:

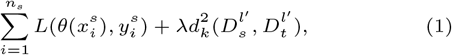

where *L* is the cross-entropy loss function for the disease status classification task, 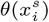 is the conditional probability that the fully connected network in Fig. 1 III assigns 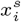 to label 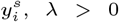 is a penalty parameter, and *l*^*′*^ is the hidden layer index where the regularizer is effective. Additional details about the domain adaptation regularizer 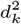 can be found in Supplementary Section 1.1. As a result, by minimizing the loss function with the MK-MMD-based adaptation regularizer, the model can learn the transferable latent features between data from different domains.

After training the deep adaptation network, we applied the trained model to convert input microbial compositional features to latent features from the fully connected network. The equation for the conversion is listed in equation 2.

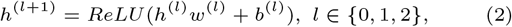

where *h*^(*l*)^ is the latent features captured from the *l*th hidden layer of the model, and *h*^(0)^ = *x. w*^(*l*)^ and *b*^(*l*)^ are the learnable weights and biases of the hidden layers. ReLU is the activation function. Finally, each input sample has corresponding latent features *h*^(2)^ that contain robust statistic patterns from compositional abundance data.

### Edge construction

Given the latent features of all samples, a sample-sample Euclidean distance matrix can be calculated. This matrix effectively captures the microbial compositional similarity across all samples. Then, we employ the *k*-nearest neighbor algorithm to determine whether two samples have a connection. In our algorithm, we will connect the *k* closest samples for each sample according to the distance matrix.

### The GCN model

After constructing the graph, we train a GCN model to classify disease status for input samples. The most important component of GCN is the graph convolution layer, which can take advantage of the topological structure for feature learning. Because both labeled and unlabeled samples are connected in the graph, GCN can also utilize node features from unlabeled nodes. If there are *n* samples in the graph, the graph convolution layer is defined as:

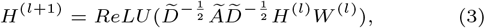

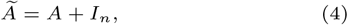

where *A* is the ℝ^*n×n*^ adjacency matrix of the inter-host microbiome similarity graph, *I*_*n*_ is an ℝ^*n×n*^ identity matrix, and *D* is the ℝ^*n×n*^ degree matrix of *Ã. H*^(*l*)^ refers to the output of *l*th hidden layer and *H*^(0)^ is the node feature matrix. *W* ^(*l*)^ represents the weight matrix of the hidden layers. As shown in equation 3, in each convolution layer, the model considers the 1-hop neighborhood of the nodes to calculate new node features. Then, the successive convolution will be applied in the *l* layers, and thus, the model can learn features based on the topological structure to enlarge the receptive field from unlabeled nodes and make a classification. All the nodes will take part in the convolution as shown in equation 3. But only the labeled nodes are used in optimizing the loss function, which is a cross-entropy loss function for the disease status classification. Finally, the model will assign disease status to unlabeled samples in the classification step.

### Discover biomarkers and analyze their contribution to the host disease status with GDmicro

In microbiome-related studies, biomarkers typically refer to species that are highly correlated with diseases and are often considered signatures of certain health conditions [4]. Biomarker discovery is an essential task that can help reveal the underlying biological mechanisms of diseases. However, identifying biomarkers using deep learning models remains challenging. Although methods such as the ablation approach are available for interpreting deep learning models [23], they cannot be employed in this study due to computational efficiency and performance issues. Specifically, the sheer number of features (over 800 in CRC datasets) makes these methods computationally expensive. In addition, in our specific case, methods like the ablation approach, which requires the removal of one feature from hundreds, cannot produce noticeable differences to the model, making feature selection highly difficult. To tackle these issues, we propose a straightforward yet highly effective performance-based method that utilizes both the input data and the GCN model to extract significant species. Furthermore, based on the identified biomarkers, we design a function that utilizes the GCN model to analyze the biomarkers’ contributions to the host disease status. We will discuss these two methods in the following paragraphs.

First, we utilize the node features in the GCN model for biomarker discovery. Once the graph is given, the performance of the GCN can vary when feeding it with different species. Therefore, we run GDmicro multiple times using a single species as the node feature each time. As a result, each species obtains a corresponding AUC on the training data, allowing for species ranking according to their respective AUCs. Higher-ranked species can be crucial biomarkers for classifying host disease status.

Second, given the identified biomarkers, we will further characterize the contribution of these biomarkers to the hosts’ disease status. To solve this problem, we first define four kinds of contributions of biomarkers to the disease status: “Increase2Disease”, “Increase2Health”, “Decrease2Disease”, and “Decrease2Health”. These contributions reflect that modifying the abundance of certain species, either through increase or decrease, may result in a sample being more likely classified as diseased or healthy, respectively. For example, “Increase2Disease” represents that increasing the abundance of the species will result in a higher probability of the sample being classified as a diseased one. Thus, we can analyze a biomarker’s contribution by changing its abundance and checking the class (health or disease) probability predicted by the model. Based on the idea, we design a function utilizing the output probability of the GCN model to calculate the score for each kind of contribution. The higher the score, the more likely the biomarker has that specific contribution to the host’s disease status. Fig. 2 illustrates the workflow of this function. If there are *n* correctly predicted samples in the training data, the contribution score of contribution type *i* (*i ∈ {*Increase2Disease, Increase2Health, Decrease2Disease, Decrease2Health*}*) for biomarker *j* can be represented as 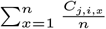, where *C*_*j,i,x*_ is the contribution variable shown in Fig. 2. More details about this function can be found in Supplementary Section 1.2.

**Fig. 2.**
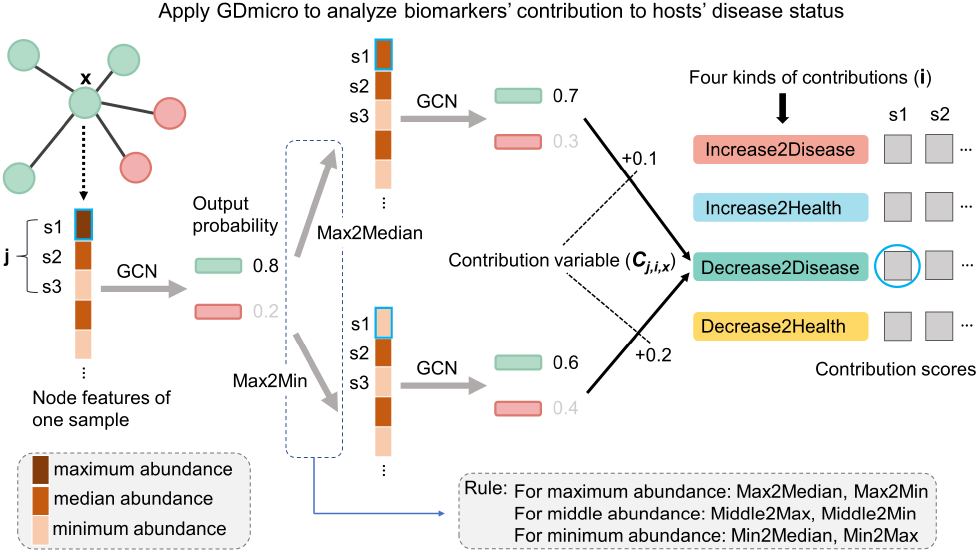
One example of using GDmicro to analyze a biomarker’s contribution to the host disease status. “s1, s2, s3”: identified biomarkers. “Rule”: the rule to change the abundance of the analyzed biomarker. For example, “Max2Median” means the abundance of the biomarker will change from the maximum value to the median value. “Contribution variable”: the absolute value of the difference between the originally predicted probability of being healthy and the predicted probability of being healthy after changing the abundance.

## Results

In this section, we describe GDmicro’s results on 10-fold cross-validation and cross-study experiments in 10 cohorts with 5 diseases. Then, we analyzed the effect of different model architectures and parameters on the performance of GDmicro. Finally, we explored the ability of GDmicro to identify biomarkers for CRC and IBD cohorts and explore their influence on hosts’ disease status. We benchmarked four state-of-art tools, including SIAMCAT [8], MetAML [7], DeepMicro [9], PopPhy-CNN [10] with GDmicro in the 10-fold cross-validation and cross-study experiments. We take microbial compositional data as input for all of these tools. All tools are run with the recommended parameters or the best performance parameters reported in the corresponding paper.

### Datasets and evaluation metrics

To benchmark with other tools [7, 9, 10, 24], we first used the same set of 5 cohorts covering 5 diseases: liver cirrhosis (Cirrhosis), colorectal cancer (CRC), inflammatory bowel disease (IBD), obesity (Obesity), and type 2 diabetes (T2D). We additionally collected 4 CRC cohorts and 1 IBD cohort for the cross-study experiment. These cohorts contain 1644 samples in total and have composition profiles available, which are generated by Metaphlan2 [25] or mOTU2 [26]. We downloaded five CRC cohorts from [4] and all other cohorts from curatedMetagenomicData package [27]. The detailed information of all cohorts is summarized in Table 1.

**Table 1.**
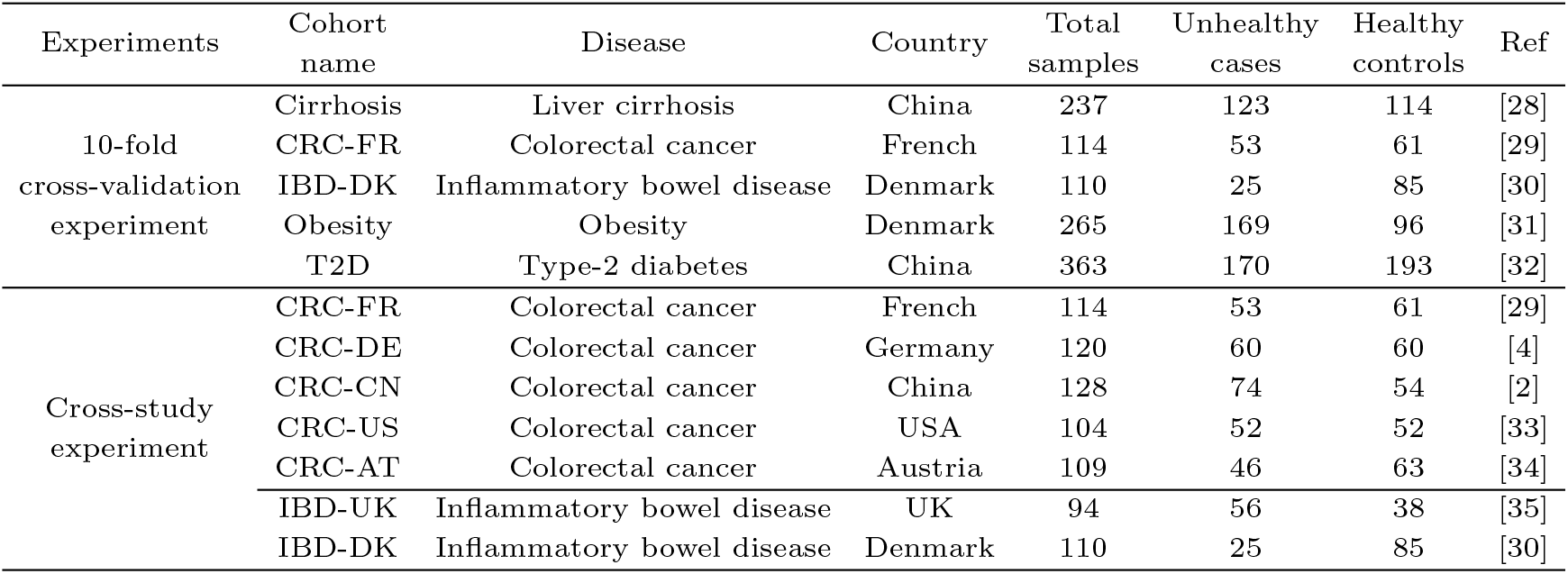
The detailed information of 10 cohorts used in this work. The first five cohorts are used for the 10-fold cross-validation experiment. CRC-FR, IBD-DK, and the remaining five cohorts are used for the cross-study experiment.

| Experiments | Cohort name | Disease | Country | Total samples | Unhealthy cases | Healthy controls | Ref |
| --- | --- | --- | --- | --- | --- | --- | --- |
| 10-fold cross-validation experiment | Cirrhosis | Liver cirrhosis | China | 237 | 123 | 114 | [28] |
|  | CRC-FR | Colorectal cancer | French | 114 | 53 | 61 | [29] |
|  | IBD-DK | Inflammatory bowel disease | Denmark | 110 | 25 | 85 | [30] |
|  | Obesity | Obesity | Denmark | 265 | 169 | 96 | [31] |
|  | T2D | Type-2 diabetes | China | 363 | 170 | 193 | [32] |
| Cross-study experiment | CRC-FR | Colorectal cancer | French | 114 | 53 | 61 | [29] |
|  | CRC-DE | Colorectal cancer | Germany | 120 | 60 | 60 | [4] |
|  | CRC-CN | Colorectal cancer | China | 128 | 74 | 54 | [2] |
|  | CRC-US | Colorectal cancer | USA | 104 | 52 | 52 | [33] |
|  | CRC-AT | Colorectal cancer | Austria | 109 | 46 | 63 | [34] |
|  | IBD-UK | Inflammatory bowel disease | UK | 94 | 56 | 38 | [35] |
|  | IBD-DK | Inflammatory bowel disease | Denmark | 110 | 25 | 85 | [30] |

Like many previous studies [7, 36, 37, 38], we selected the area under the curve (AUC) as the evaluation metric in this work, which summarizes the true positive and false positive rates and has a robust evaluation for the unequal ratio of each outcome. For the 10-fold cross-validation experiment, each experiment is repeated 10 times. Then, the margin of error for the mean of all experiments is calculated with a 95% confidence interval, and it is defined as:

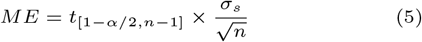

where 1 *− α* is the significance level, *n* is the sample size, *t*_[1*−α/*2,*n−*1]_ refers to critical value of t-distribution with degrees of freedom n-1 for an *α/*2 area of for the upper tail, and *σ*_*s*_ is the sample standard deviation.

### 10-fold cross-validation experiment

To evaluate the performance of GDmicro, we applied all tools to classify disease status for samples from five cohorts (upper part in Table 1). We conducted 10-fold cross-validation using StratifiedKFold (k=10) function in the sklearn package. The mean AUC and the margin or errors of all tools were recorded in Table 2. GDmicro achieved the best performance in all tested cohorts. In particular, GDmicro achieved 0.923 and 0.936 AUC in the CRC-FR and IBD-DK cohorts, which are 6.4% and 6.3% improvements over the second-ranked tool, respectively. Although some tools have competitive results, they have larger fluctuations than GDmicro. For example, while SIAMCAT achieved 0.949 AUC in the Cirrhosis cohort, its mean AUC in the T2D cohort was only 0.691. Similarly, DeepMicro had 0.940 AUC in the Cirrhosis cohort, but its mean AUC in the CRC-FR cohort was only 0.803, which was much lower than the top two tools. The results also reveal that the performance of all methods was not satisfactory in the Obesity and T2D cohorts. A possible reason is that the microbial features do not have strong correlations with the disease as reported in the previous study [39]. Overall, GDmicro outperformed all other tools in the five tested cohorts.

**Table 2.**
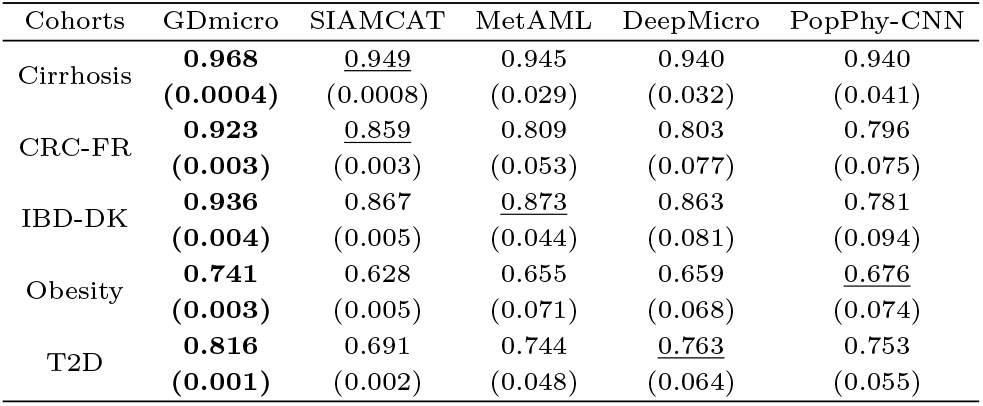
The mean AUC (10-fold cross-validation) of 5 tools on five popular disease cohorts. The values in parenthesis refer to the margin of errors. Bold: the best performance of each cohort. Underlined: the second-best performance of each cohort.

### Cross-study experiment

Test data in real-world applications usually come from different studies or domains from the training data. Thus, it is imperative to evaluate the robustness of the model across different domains. Because CRC and IBD are more associated with human gut microbes based on previous literature [40, 41] and our experiments, we conducted cross-study experiments for these diseases.

We collected additional human gut microbiome data for CRC and IBD. The additional four CRC cohorts were all sampled from different individuals, while some samples in the additional IBD cohort were sampled from the same individual at different ages. In total, we have five CRC cohorts and two IBD cohorts. All these cohorts are from different countries. Then, we used the leave-one-study-out (LOSO) strategy to evaluate the performance of different tools. Specifically, we remove all samples from one study in the training data and retrain the model. Then the model is tested on the removed samples. This setup mimics the real scenario where the query is from a different research group. The AUC of all tools in seven test cohorts was shown in Fig. 3.

**Fig. 3.**
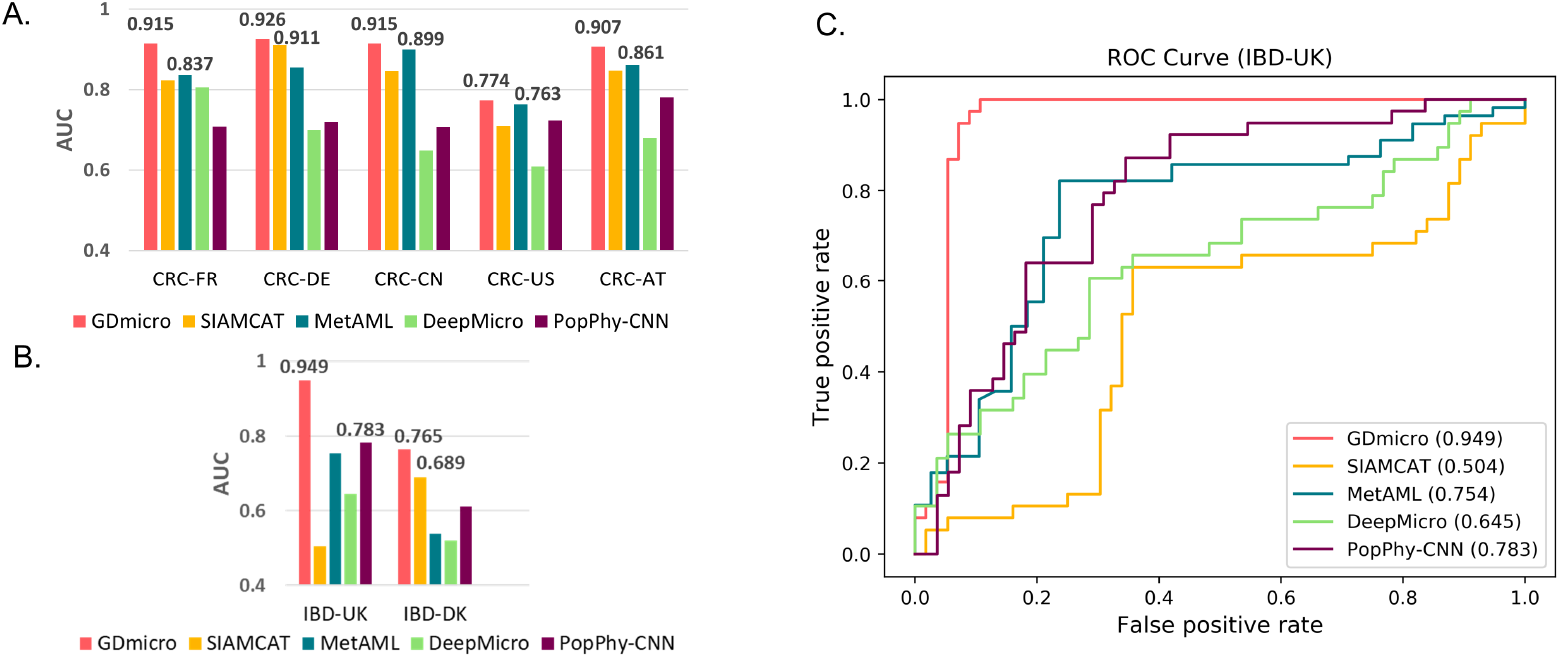
(A-B). The cross-study AUC of five tools in LOSO experiments of CRC (A) and IBD (B). The X-axis represents the test data in each LOSO experiment. For example, CRC-FR represents we train the model using CRC cohorts from four other countries and test on the CRC cohort from French. The values on the bar refer to the best and second-best AUC. The detailed AUC values of each tool are recorded in Supplementary Table S1. (C). ROC curves in the IBD-UK cohort. The value in the parentheses represents AUC for each tool.

Considering that both CRC-FR and IBD-DK cohorts were used in the 10-fold cross-validation experiment (Table 1), we first analyzed the performance change of different tools in these two cohorts. GDmicro, MetAML, and DeepMicro achieve comparable performance to the 10-fold cross-validation experiment in the CRC-FR cohort, while the AUC of SIAMCAT and PopPhy decreases more than 3%. However, the AUC of all tools decreases in the IBD-DK cohort. This result indicates that some models can still maintain good robustness in the cross-study experiments when sufficient training samples are available (e.g., 421 training samples for CRC-FR). However, when the training sample size is small, and some samples are from the same individual (e.g., 94 training samples from 50 individuals for IBD-DK), the classification of cross-study samples can be a challenge for all tested tools. Then we will discuss the overall performance of these tested tools in all seven test cohorts.

As shown in Fig. 3A and 3B, GDmicro achieved the best and most stable performance in the LOSO experiments. Compared to the second-best result, GDmicro improved the AUC from 0.837 to 0.915 and 0.861 to 0.907 in CRC-FR and CRC-AT cohorts, respectively. MetAML and SIAMCAT achieved comparable performance in the CRC-US cohorts. However, their performance was not stable in the remaining cohorts. Although DeepMicro and PopPhy-CNN achieved satisfactory performance in the previous experiments, their performance was poor in most tested cohorts. In the two IBD cohorts, GDmicro achieved more than 15% improvement in AUC in the IBD-UK cohort. Thus, we further draw a ROC curve of each tool in the IBD-UK cohort in Fig. 3C, which shows that GDmicro has more reliable classification results than other tools. However, all the tools do not perform well in the IBD-DK cohort, which contains data from the fewest number of individuals out of the seven cohorts. This result indicates that the performance of the learning-based methods may fluctuate with the change of the training cohort size. Nevertheless, GDmicro achieves more than 5% improvement in AUC.

From Fig. 3A and 3B, we also observed that machine-learning-based tools (MetAML and SIAMCAT) tend to have better performance than deep-learning-based tools (DeepMicro and PopPhy-CNN) in most LOSO experiments. This result indicated that deep-learning-based tools are more likely to overfit the training set because of the complexity of the model. However, by using a deep adaptation network to learn the transferable latent features, GDmicro maintains good generalization capabilities in the cross-study host disease status classification task.

### Ablation study and parameter analysis

In this experiment, we study how different architectures and parameters influence the performance of GDmicro using ablation study and parameter analysis. Specifically, we analyzed the influence of the adaptation loss function, GCN model, and hyper-parameter *k* in the *k* NN graph on the performance of GDmicro. To be more consistent with the usage of real-world data, we analyzed datasets of the cross-study experiment. The analysis results show that the current architecture outperforms all other tested architectures, demonstrating the efficiency of domain adaptation and the GCN model (Supplementary Figure S1A). In addition, we also found that the performance of GDmicro is not very sensitive to *k* (Supplementary Figure S1B). By default, we use *k* = 5 to construct the *k*NN graph. Additional details regarding this experiment can be found in Supplementary Section 2.1.

### GDmicro identifies important biomarkers and analyzes their contribution to CRC and IBD

In this experiment, we applied GDmicro to identify important biomarkers and explore their contribution to hosts’ disease status from cohorts used in the cross-study experiment. As mentioned in the Methods section, GDmicro could identify important biomarkers using a performance-based method. According to the method, the species with better classification ability will have higher AUC. Thus, we can evaluate the classification ability of all species in the given data and record their AUC. Then, we selected the top 10 species with the highest AUC values for further analysis. The identified biomarkers and their relative abundance distribution are shown in Supplementary Figure S2. We observed that many identified species of CRC cohorts were enriched in CRC samples, and their relative abundance was also higher. Unlike CRC, many identified species in the IBD cohort were enriched in healthy samples, which was in line with previous observations [42, 43]. Given that abundance-based statistical analysis is a prevalent method for discovering microbial biomarkers, we examined the differences between biomarkers identified by GDmicro and those detected using the Wilcoxon test, a popular statistics-based approach. The details about using the Wilcoxon test to discover biomarkers are described in Supplementary Section 2.2. In the following paragraphs, we first analyzed the biomarkers identified by GDmicro and the Wilcoxon test in terms of robustness, discriminatory capabilities, and cross-study consistency. Then, we investigated the contribution of these cross-study biomarkers to the host disease status.

To assess the robustness of identified biomarkers, we further calculated the p-value of identified features in each study (Supplementary Figure S3) and defined two kinds of features. The first feature is called “robust feature” that exhibits a consistent positive or negative p-value of less than 0.05 across all studies. The second type of feature, called “inconsistent feature”, refers to features that exhibit both positive and negative p-values across different datasets. In real-world applications, we expect that effective biomarker discovery methods can uncover more robust features while reducing the number of inconsistent features. As Fig. 4A shows, GDmicro identified more robust features and produced fewer inconsistent features than the statistics-based method.

**Fig. 4.**
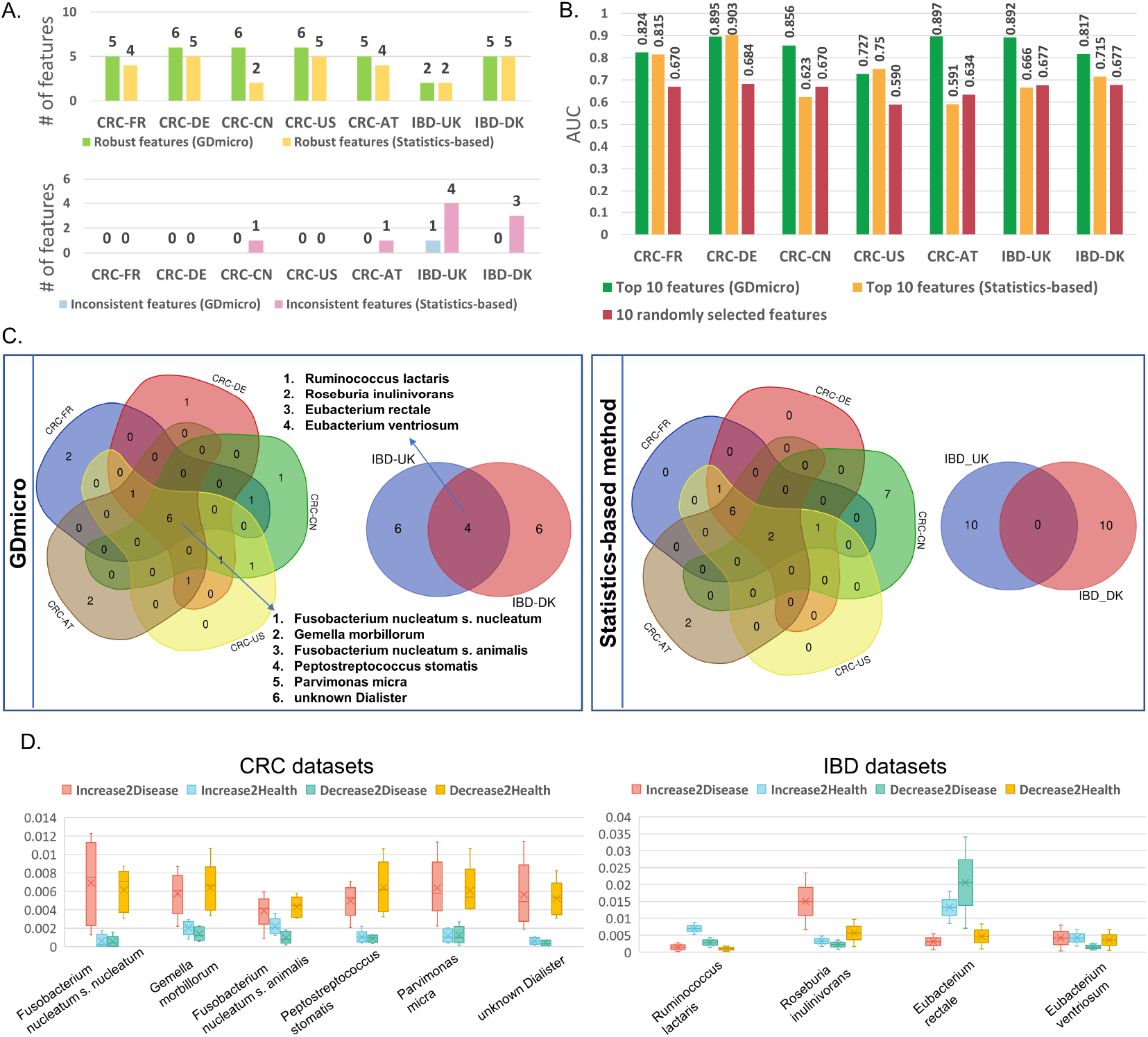
(A). The number of robust and inconsistent features identified by GDmicro and the statistics-based method (Wilcoxon test). (B). AUC comparison across studies in the LOSO experiment using the top 10 features identified by three different methods. The AUC of “10 randomly selected features” is the average AUC of 5-time repeated experiments. (C). Venn diagrams for the identified top 10 biomarkers in different cohorts by GDmicro and the statistics-based method. (D). The boxplot of contribution scores of cross-study biomarkers in CRC and IBD datasets. The Y-axis represents the contritbution score values of the biomarker in five CRC and two IBD cohorts.

To evaluate the discriminatory capabilities of the identified biomarkers, we conducted the LOSO experiment using only the top 10 species selected by GDmicro and the Wilcoxon test, respectively. For comparison, we randomly selected 10 features from each study and reran the LOSO experiment, repeating this five times to avoid data bias. As shown in Fig. 4B, GDmicro consistently achieved higher AUC values in most tested datasets than the Wilcoxon test and outperformed the set with randomly selected species in all datasets. This result shows that the top-ranked species identified by GDmicro have better discriminatory capabilities, and an additional experiment further confirmed this superiority when using the top 50 selected features (Supplementary Section 2.2 and Figure S4).

Previous studies have reported that biomarkers with cross-study consistency (aka cross-study biomarkers) are crucial for disease identification and are valuable for clinical diagnosis [4, 44]. Thus, we compared the top 10 biomarkers identified by GDmicro and the Wilcoxon test across studies. Figure 4C shows that GDmicro identified six cross-study biomarkers for CRC and four for IBD, while the Wilcoxon test detected only two for CRC. Many cross-study biomarkers identified by GDmicro are highly associated with CRC and IBD [1, 29, 45]. For example, *Gemella morbillorum* and *Parvimonas micra* are reported as biomarkers for non-invasive diagnosis of CRC [46], while *Eubacterium rectale*, a beneficial bacteria, decreases in IBD patients [47]. To know how these cross-study biomarkers affect hosts’ disease status, we plotted their contribution scores, as calculated by GDmicro, in Figure 4D. The result shows that all CRC biomarkers may promote CRC, consistent with previous observations [4]. Conversely, IBD biomarkers play different roles: *Ruminococcus lactaris* and *Eubacterium rectale* tend to inhibit IBD, while the remaining two species prompt it. These results also align with the four species’ relative abundance distribution (Supplementary Figure S1).

## Discussion

In this work, we demonstrate that GDmicro outperforms the state-of-art tools in classifying host disease status. To reduce the domain discrepancy between data, we use a deep adaptation network to learn the transferable latent features from compositional abundance data. Then, we build an inter-host microbiome similarity graph and apply a GCN model to integrate structural and abundance features and utilize information from unlabeled and limited labeled samples. As a result, GDmicro can achieve more accurate and robust host disease status classification. Furthermore, GDmicro demonstrates superior accuracy in identifying disease-related species compared to the statistics-based method, and it elucidates their contribution to the host’s disease status. This insight offers valuable information for biomarker discovery.

Although GDmicro has improved host disease status classification, we have two goals to optimize in future work. First, current efforts are aimed at classifying whether a host has a particular disease or not. However, in practical applications, input samples may suffer from multiple diseases. Therefore, we intend to combine the graphs of different diseases for GCN to explore whether GDmicro can classify multiple diseases at the same time. Second, we will consider extending GDmicro to accommodate temporal longitudinal microbiome data. To support such an extension, methods like Long Short-Term Memory Networks will be integrated into our current architecture for feature extraction and classification.

## Data Availability

The source code of GDmicro and all datasets used in this work are freely available at https://github.com/liaoherui/GDmicro.

