## Supplementary file 1 for "GDmicro: classifying host disease status with GCN and Deep adaptation network based on the human gut microbiome data"

June 13, 2023

### 1 Supplementary Methods

#### 1.1 MK-MMD-based adaptation regularizer

In equation (1) of the main article, $d_k^2$ is the squared formulation of MK-MMD. Suppose the source domain and target domain are characterized by probability distributions $p$ and $q$, then the $d_k^2$ is defined as:

$$d_k^2(D_s, D_t) \triangleq \|E_p[\phi(x^s)] - E_q[\phi(x^t)]\|^2_{\mathcal{H}_k}, \quad (1)$$

$$\text{where } E_{x\sim p} f(x) = \langle f(x), \mu_k(p)\rangle_{\mathcal{H}_k}, \quad (2)$$

where $\phi$ is hidden layers of the network, $\mathcal{H}_k$ is the reproducing kernel Hilbert space endowed with a characteristic kernel $k$, and $\mu_k(p)$ is the mean embedding of distribution $p$ in $\mathcal{H}_k$. The main purpose of MK-MMD is to minimize the $d_k^2$ such that the two domain distribution $p$ and $q$ become closer. When $d_k^2 = 0$, $p$ will be equal to $q$. Thus, the MK-MMD is able to reflect the discrepancy between the source and target. As a result, by minimizing the loss function with the MK-MMD-based adaptation regularizer, the model can learn the transferable latent features between data from different domains.

#### 1.2 The calculation of the contribution score of biomarkers

As outlined in the main article, to determine the contribution of biomarkers to the host's disease status, we first establish four types of biomarker contributions: "Increase2Disease", "Increase2Health", "Decrease2Disease", and "Decrease2Health". With these four contribution types defined, we design a function to calculate the corresponding score for each type based on the Graph Convolutional Network (GCN) model.

Specifically, as shown in Fig. 2 of the main article, we begin by recording the classification probability for each sample. Subsequently, we modify the abundance of an identified biomarker in a sample and document the new classification probability. Six modification rules are applied: "Max2Median", "Max2Min", "Median2Max", "Median2Min", "Min2Max", and "Min2Median". Each rule represents the method of altering the abundance of the identified biomarker. For example, "Max2Median" indicates that the abundance of the selected biomarker is the maximum value among all species in the given sample, and will be adjusted from the maximum to the median value.

With the original classification probabilities and those from various change rules, we can compute the contribution variable for each biomarker. This variable is the absolute value of the difference between the originally predicted probability of being healthy and the predicted probability of being healthy after altering the abundance. This value reflects the biomarker's contribution to disease status in the given sample.

Lastly, we calculate the contribution variables of the identified biomarkers for all samples. Based on these values, we determine the contribution score of specific biomarkers to disease status using the formula discussed in the main article. The formula is $\sum_{x=1}^{n} \frac{C_{j,i,x}}{n}$, where $n$ represents correctly predicted samples in the training data, $x$ is the specific sample, $i$ is the specific contribution type, $j$ is the identified marker, and $C_{j,i,x}$ is the contribution variable.

### 2 Supplementary Experiments

#### 2.1 Ablation study and parameter analysis

In this experiment, we study how different architectures and parameters influence the performance of GDmicro using ablation study and parameter analysis. Specifically, we analyzed the influence of the adaptation loss function, GCN model, and hyper-parameter $k$ in the $k$NN graph on the performance of GDmicro. To be more consistent with the usage of real-world data, we analyzed datasets of the cross-study experiment.

As discussed in the Methods section, the loss function used in the deep adaptation network is based on multiple kernel variants of maximum mean discrepancies (MK-MMD), which aims to reduce domain discrepancy between data from different studies. To show the effect of different loss functions on the performance of GDmicro, we repeated the analysis in the cross-study experiment with and without MK-MMD-based loss. When the loss function only contains the cross-entropy loss, the model is a multi-layer fully connected network (aka multi-layer perceptron or MLP) that ignores the domain discrepancy. In addition, to know whether the GCN model improved the classification performance, we also combined the deep adaptation network and MLP for host disease status classification in the cross-study experiment.

As shown in Supplementary Figure S1A, GDmicro with default architecture achieved better performance than the model without domain adaptation in five out of seven tested datasets. This result indicates that the MK-MMD-based loss improves the model's robustness by learning transferable latent features. Related to this, the deep adaptation network outperformed MLP in all tested cohorts, which demonstrated the deep adaptation network improved the classification robustness by minimizing the domain discrepancy. We also noticed that GDmicro with default architecture achieved better performance than the single deep adaptation network and MLP, demonstrating that the GCN model improved the classification AUC by incorporating structural and compositional abundance features and utilizing information from unlabeled samples.

The hyper-parameter $k$ is an important parameter for the $k$NN graph, which determines the graph's topological structure. Thus, we investigated the performance of GDmicro under different $k$ by repeating the analysis in the cross-study experiment with $k \in \{3, 5, 7, 10\}$. Supplementary Figure S1B shows the performance of GDmicro when $k$ varies from 3 to 10. As shown in Supplementary Figure S1B, the performance of GDmicro doesn't fluctuate much in all tested datasets with the change of $k$, which indicates that GDmicro is not very sensitive to $k$. By default, we use $k = 5$ to construct the $k$NN graph.

#### 2.2 LOSO experiments with top 50 features selected by different methods

To identify biomarkers with the Wilcoxon test, we calculated the p-value of each feature using all the training data. In this experiment, a positive p-value signifies that the feature is enriched in disease samples, whereas a negative value indicates enrichment in healthy samples. Subsequently, all features are sorted from smallest to largest based on the absolute value of their p-values. To avoid data bias, we repeated the LOSO experiment with the top 50 features identified by GDmicro and the statistics-based method. The result shows that the average AUC for GDmicro is 0.891, while the average AUC for the statistics-based method is 0.869 (Supplementary Figure S4), a finding consistent with the result observed for the top 10 features.

### 3 Supplementary Figures

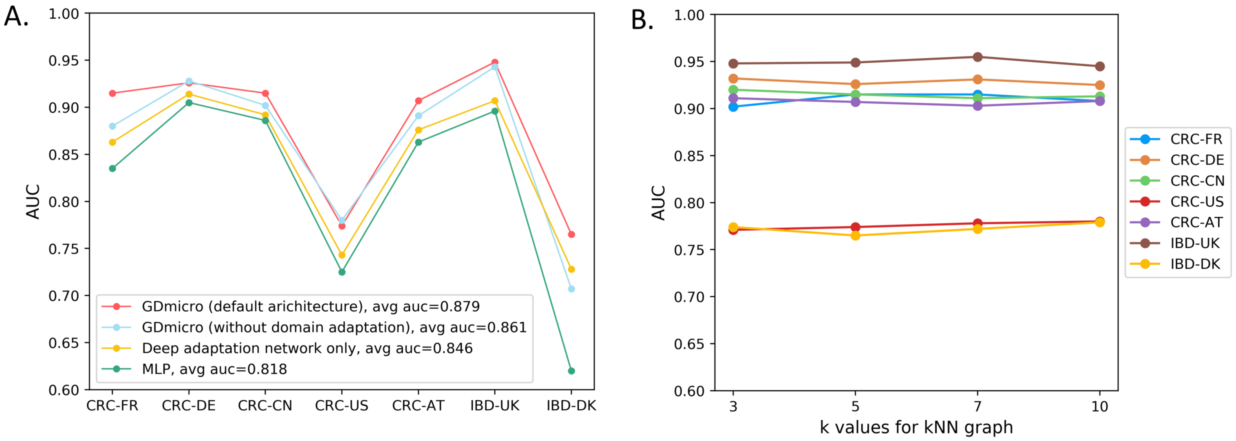


**Supplementary Figure S1.** (A). The AUC of deep adaptation network, MLP, and GDmicro with different loss functions in LOSO experiments. "avg auc": the average AUC of the tested model in seven cohorts. Deep adaptation network only: applying a single deep adaptation network (Fig. 1 III in main article) to classify host disease status with species abundance data. MLP: applying a multi-layer perceptron to classify host disease status with species abundance data. (B). The AUC of GDmicro with different *k* values in LOSO experiments.

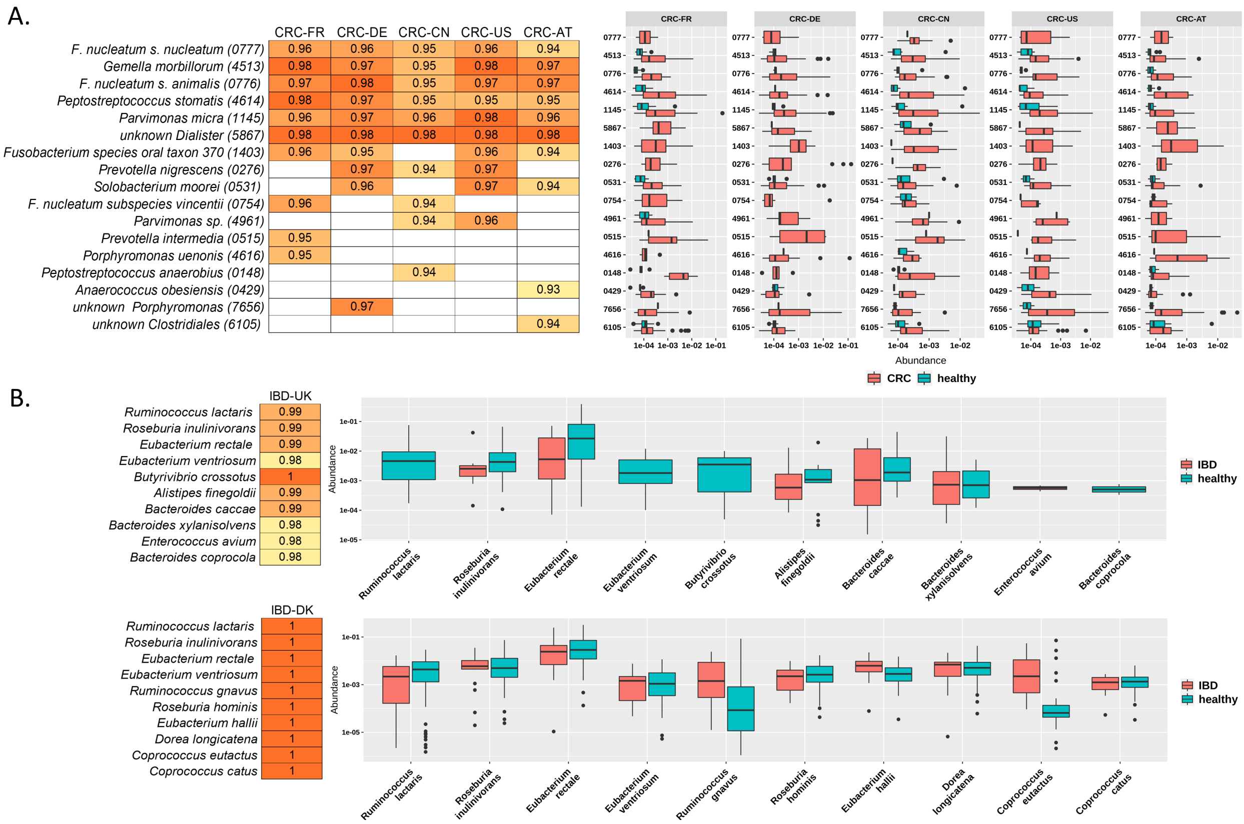


**Supplementary Figure S2.** (A). The top-10 disease-related species identified by GDmicro on 5 CRC datasets. The values in the cell represent the AUC using that species as the node feature. The boxplot shows the abundance distribution of identified species in CRC patients and healthy samples. (B). The top-10 disease-related species identified by GDmicro on 2 IBD datasets. The values in the cell represent the AUC using that species as the node feature. The boxplot shows the abundance distribution of identified species in IBD patients and healthy samples.

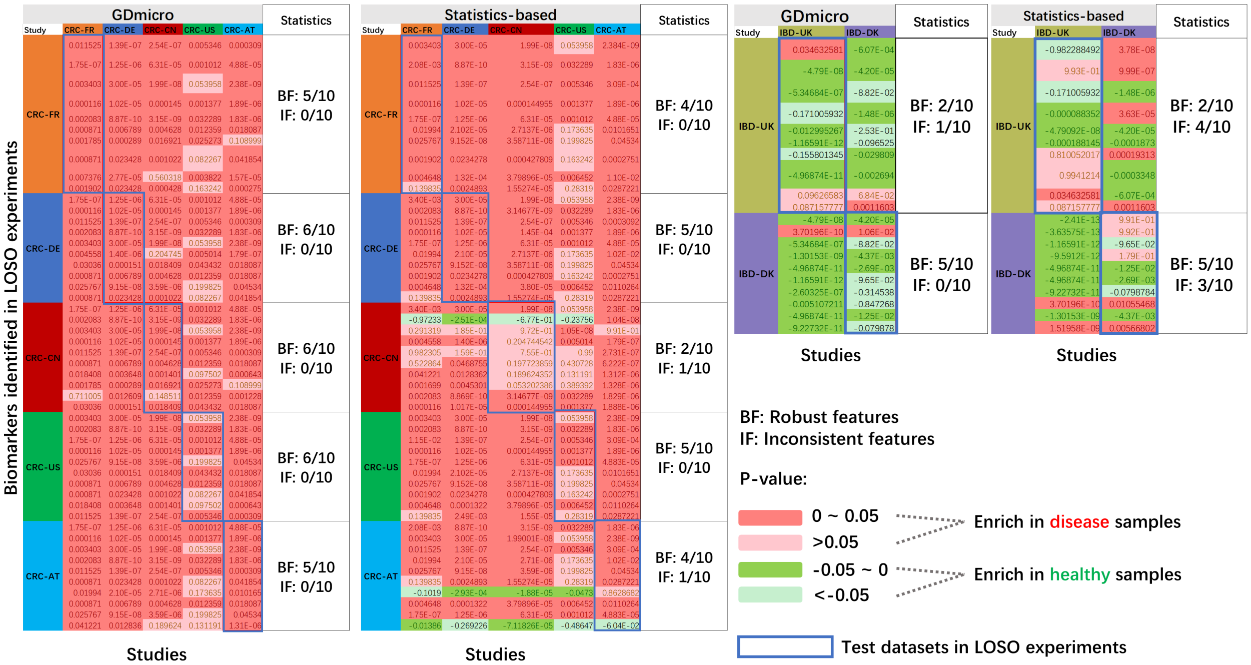


**Supplementary Figure S3.** P-values of top-10 features identified by GDmicro and statistics-based method in CRC and IBD datasets. Each row represents a feature identified in the test dataset of the LOSO experiment, and each cell corresponds to a p-value of the feature in the specific study. The p-value is calculated using the Wilcoxon test, where a positive value (marked as red) indicates that the feature is enriched in disease samples, while a negative value (marked as green) signifies enrichment in healthy samples. Robust features: these are features exhibiting a consistent positive or negative p-value of less than 0.05 across all tested datasets. Inconsistent features: these are features that display both positive and negative p-values within the tested datasets.

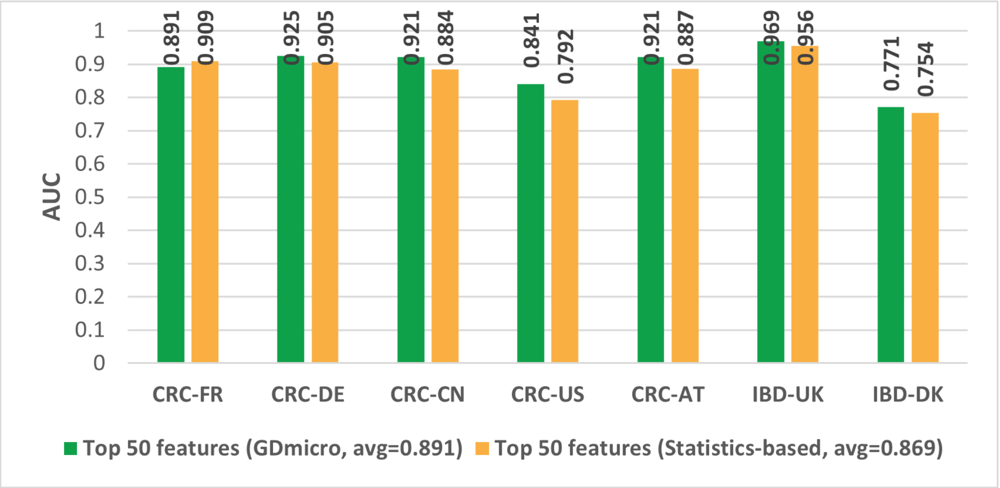


**Supplementary Figure S4.** The AUC of using the top 50 biomarkers identified by GDmicro and the Wilcoxon test in the LOSO experiment. “avg auc”: the average AUC of the tested method in seven cohorts.

### 4 Supplementary Tables

| Datasets | GDmicro | SIAMCAT | MetAML | DeepMicro | PopPhy-CNN |
|---|---|---|---|---|---|
| CRC-FR | **0.915** | 0.823 | 0.837 | 0.806 | 0.708 |
| CRC-DE | **0.926** | 0.911 | 0.855 | 0.699 | 0.72 |
| CRC-CN | **0.915** | 0.846 | 0.899 | 0.649 | 0.707 |
| CRC-US | **0.774** | 0.71 | 0.763 | 0.609 | 0.723 |
| CRC-AT | **0.907** | 0.847 | 0.861 | 0.68 | 0.781 |
| IBD-UK | **0.949** | 0.504 | 0.754 | 0.645 | 0.783 |
| IBD-DK | **0.765** | 0.689 | 0.539 | 0.52 | 0.611 |

**Supplementary Table S1.** The cross-study AUC of five tools on seven test cohorts in the LOSO experiment. Bold: the best performance of each cohort.
